# Self-supervised machine learning methods for protein design improve sampling, but not the identification of high-fitness variants

**DOI:** 10.1101/2024.06.20.599843

**Authors:** Moritz Ertelt, Rocco Moretti, Jens Meiler, Clara T. Schoeder

**Affiliations:** Institute for Drug Discovery, Leipzig University Faculty of Medicine, Leipzig, Germany; Center for Scalable Data Analytics and Artificial Intelligence ScaDS.AI, Dresden/Leipzig, Germany; Department of Chemistry, Vanderbilt University, Nashville, Tennessee, United States of America; Center for Structural Biology, Vanderbilt University, Nashville, Tennessee, United States of America

## Abstract

Machine learning (ML) is changing the world of computational protein design, with data-driven methods surpassing biophysical-based methods in experimental success rates. However, they are most often reported as case studies, lack integration and standardization across platforms, and are therefore hard to objectively compare. In this study, we established a streamlined and diverse toolbox for methods that predict amino acid probabilities inside the Rosetta software framework that allows for the side-by-side comparison of these models. Subsequently, existing protein fitness landscapes were used to benchmark novel self-supervised machine learning methods in realistic protein design settings. We focused on the traditional problems of protein sequence design: sampling and scoring. A major finding of our study is that novel ML approaches are better at purging the sampling space from deleterious mutations. Nevertheless, scoring resulting mutations without model fine-tuning showed no clear improvement over scoring with Rosetta. This study fills an important gap in the field and allows for the first time a comprehensive head-to-head comparison of different ML and biophysical methods. We conclude that ML currently acts as a complement to, rather than a replacement for, biophysical methods in protein design.

## Introduction

Computational design and engineering of proteins has been a long-standing goal of the scientific community, allowing for the rapid generation of new protein drugs and materials. Traditionally, software suites for molecular modeling and design such as Rosetta^1^ have been used and developed for the design of the first *de novo* protein^2^, enzymes^3, 4^, and antibodies^5^, amongst other examples. These successes were made possible not only by the sequence mutagenesis and design tools included in Rosetta but through the efficient combination with other protocols, including protein structure prediction, symmetry, filtering, modeling, refinement, and docking. Additionally, the establishment of the RosettaScripts^6, 7^ framework and PyRosetta^8^ allows for the generation of tailor-made protocols without the need to edit the C++ source code, facilitating quick protocol development for non-computational scientists. Over time the scientific community has added many improvements to these software packages to close in on the two fundamental underlying problems of protein sequence design: the sampling problem, e.g. combining packing with relaxation^9, 10^, and the scoring problem, continuously optimizing the scoring function^11^.

Recently, machine learning (ML) methods have shown impressive performance on tasks like protein structure prediction^12–14^, molecular docking^15, 16^, protein sequence design^17^ and protein engineering^18^. Especially in the area of *de novo* protein design, the deep learning-based method ProteinMPNN has led to major breakthroughs, including the design of two-component nanomaterials^19^ and the *de novo* design of luciferases^20^. In the realm of protein engineering ProteinMPNN was successfully applied to e.g. the design of myoglobin and tobacco etch virus protease variants with improved stability^21^. Similarly, protein language models (PLMs) have both created *de novo* proteins^22^ and engineered existing ones, for example increasing the binding affinity of a mature antibody by sevenfold^18^.

It remains however an outstanding question, in which regard ML models outperform classic biophysical design algorithms, which in return might be able to inform us about the true power of these models and the gaps in biophysically driven design algorithms. Additionally, the resulting complex pipelines of multiple Python packages can be prone to technical debt, lack of testing and hard to maintain^23^. Furthermore, often just the core functionality needed to reproduce publication results for a specific dataset is provided. This results in time-consuming user re-implementation tasks, starting from efficiently parsing protein structure files (which can be surprisingly non-trivial for example when other entities in addition to canonical amino acids are involved), to scoring mutations. These personally adapted protocols are hard to compare and transfer to other software set-ups, resulting in user frustration and lack of reproducibility. After embedding the evolutionary scale modeling (ESM) protein language model (PLM) family^14, 22, 24–26^ in Rosetta^27, 28^, we realized the benefits of a shared interface and testing environment using the C++ Tensorflow^29^ and LibTorch^30^ libraries, and therefore set out to streamline and expand this interface to other models.

In this work, we tested in which regard novel self-supervised ML methods outclass biophysical-based methods like Rosetta and identified best practices for new design projects. To do so, we made use of existing protein fitness landscape datasets to benchmark the new tools on common tasks like improving protein binding affinity or enzymatic activity, assessing their ability to generalize without further downstream training. Two of the main objectives of a protein engineering campaign are the generation of new candidates (sampling mutations) and then ranking these candidates (scoring mutations). Therefore, predictive models (termed “oracles”) were trained on large scale mutagenesis datasets to analyze the sampling and scoring behavior of sixteen different protocols. We found that while ML methods are significantly better at purging the sequence space from deleterious mutations, scoring and ranking the resulting candidates remains a challenge in protein design.

## Results

### Definition of appropriate benchmark sets and ML models

Inverse folding models like ProteinMPNN have been benchmarked on tasks such as sequence recovery and finally on the experimental success of their designs^17^. Similarly, pre-trained masked-language models like ESM2^14, 24^ and masked inverse folding with sequence transfer^31^ (MIF-ST) were benchmarked by assessing their ability to predict the effects of mutations in a zero-shot fashion or by fine-tuning the models on a specific downstream task with an evaluation of their predictions. However, these methods are not continuously tested on the same benchmark set. We set out to test different protocols in their ability to generate and rank novel candidates with improved attributes, without fine-tuning for the given task (**Fig. 1 A**). This setup tests the ability to generalize from pre-training alone, as the models have not been further trained on the datasets used here for validation. To accurately assess the sequences, we focused on four case studies where large-scale fitness landscapes were available. These comprehensive datasets were used to train a simple predictive model, termed “oracle”, enabling us to predict different fitness aspects *in silico* (**Fig. 1 B**). This allowed us to evaluate sixteen different protocols from various approaches. Additionally, while protein design studies commonly focus on selecting designs for experimental verification that have the best predicted success rates, here we are especially interested in the pitfalls of different methods. We chose ProteinMPNN^17^, MIF-ST^31^, and ESM2^14, 24^ to cover the spectrum of structure- and sequence-based ML models and allow for a head-to-head comparison.

**Fig. 1.**
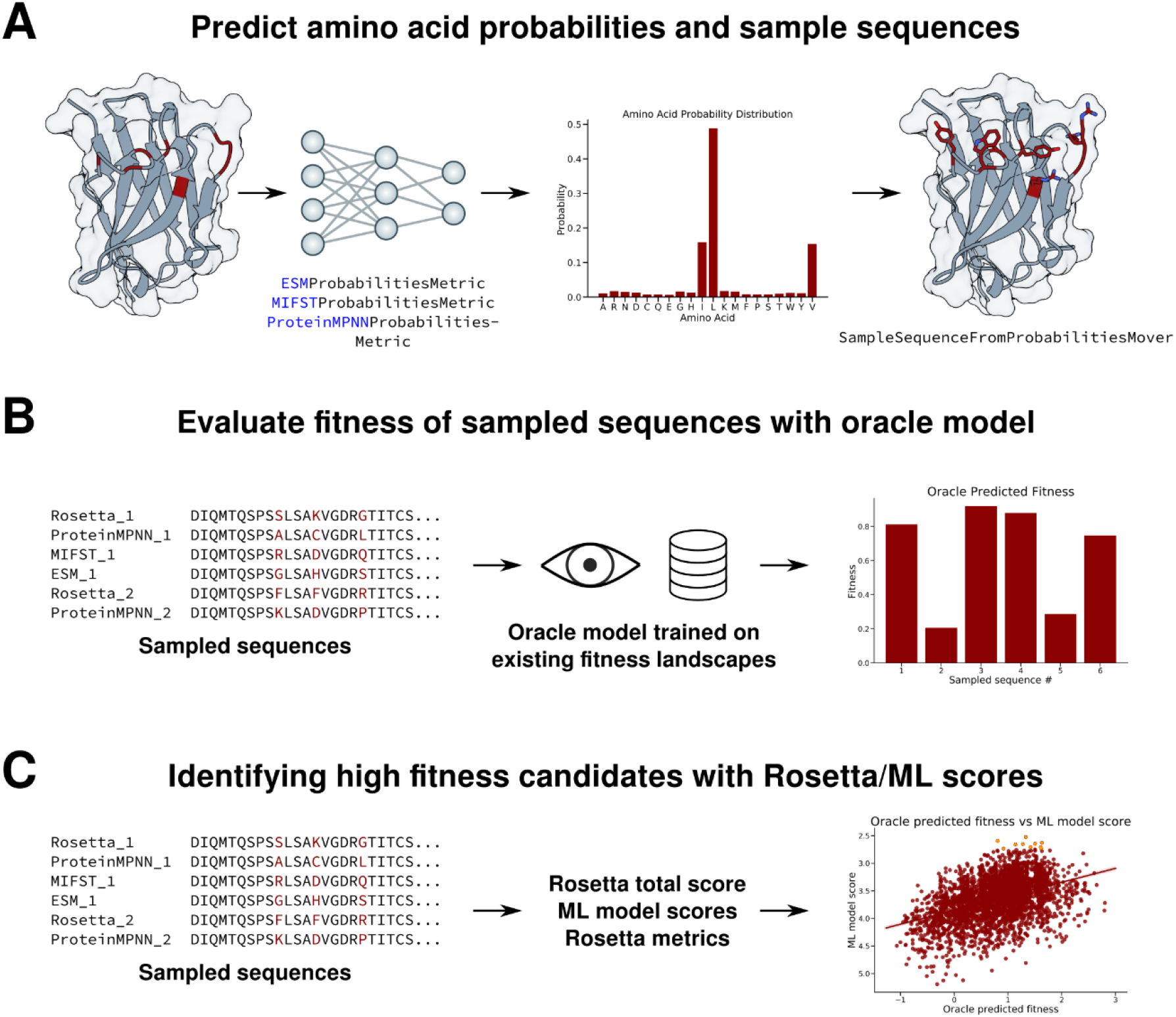
ML-support framework in in Rosetta. **A)** Implemented RosettaScripts elements include metrics to run different ML models for predicting amino acid probabilities and a mover to directly thread mutations sampled from the predictions on to the structure loaded in Rosetta. **B)** New variants sampled by different models and option settings were evaluated *in silico* through an oracle model trained on existing fitness landscapes. **C)** The sampled variants and resulting predicted fitness scores were used to evaluate the correlations and ranking behavior of different Rosetta/ML model scores and metrices.

### Creating a common interface for sampling sequences from predicted amino acid probabilities

To compare the ML approaches against Rosetta, designs were generated with the `FastDesign` mover as a baseline. We first created a general `SimpleMetric`^7^ class for holding predicted probabilities, called `PerResidueProbabilitiesMetric`. Subsequently, we created a metric for each model that can be used through the `RosettaScripts` framework, where the user can provide a `ResidueSelector` to specify subsets of the protein for prediction. Our approach to derive mutations from the predicted probabilities was implemented as `SampleSequenceFromProbabilities` mover. While sampling mutations for the full sequence of a protein might be useful for e.g., designing a *de novo* protein or the comprehensive re- design of a scaffold, many other common protein engineering tasks seek to maximize impact while minimizing the number of mutations. To cover both use cases, we first rank the to-be- designed positions by the maximum difference of probability to the current sequence, and then rank the amino acids for each position based on their predicted probability. The mover’s options allow users to adjust temperature parameters for the selection of both position and amino acid which influence the determinism of their selection, in addition to specifying the desired number of mutations. For benchmarking each implemented model, their respective `PerResidueProbabilitiesMetric` (stored predicted probabilities) was used as input for the `SampleSequenceFromProbabilities` mover, as well as an average of all three model predictions using the `AverageProbabilitiesMetric`. A detailed description of all newly implemented features can be found in the Supplementary Information. Three different temperatures (0.3, 1.0, 1.5) were set for amino acid sampling to find the optimum between exploitation and exploration of predicted probabilities. Additionally, we hypothesized that sampling conformational space during design could be beneficial, therefore a protocol was tested consisting of three rounds of ProteinMPNN design each followed by a relaxation of the structure thereby increasing backbone flexibility, as described by Bennett *et al*.^32^. Lastly, as the probabilities sampled from PLMs are conditional given the contextual sequence, the `IteratedConvergenceMover` (IC) was used to sample only one mutation at a time until the model score converged. 1000 trajectories were produced for each approach and all unique sequences were subsequently analyzed.

### Setup for identifying best practices in ranking designed sequences

After sampling, we set out to find best practices in identifying high fitness candidates, analyzing the correlation of ML scores to protein fitness and their ranking accuracy. To do so, we created a `PseudoPerplexityMetric` using the predicted probabilities of ML models to score the likeliness of a given sequence and compared these to Rosetta scores including total score, dG separated and shape complementarity. Additionally, we analyzed scores derived from AlphaFold2 (AF2) structure prediction^12^ as orthogonal approach. For the ranking accuracy we sorted all sampled sequences irrespective of design method by each score and analyzed the top ten candidates. This sort of ranking is common when choosing designs for experimental validation in a low to moderate throughput setting.

### Sampling GB1 variants with at least twofold improved fitness required higher sampling temperatures

Wu *et al*.^33^ collected a dataset compromising ∼150,000 mutations of four sites in immunoglobin-binding protein G domain B1 (GB1), a protein commonly used in the purification of antibodies. The measured fitness of the resulting variants is both influenced by the stability of the resulting protein and the binding affinity towards IgG-Fc, relative to the wild type (**Fig. 2 A, B**). Using this dataset, a ridge regression model was trained (spearman correlation of 0.79) and used to predict the fitness of designs sampled by the above described protocols, without further training of the ML models used for sampling. Additionally, we provide an analysis using only measured fitness values instead of the oracle predictions (**Supp. Fig. 2**), which showed the same behavior of different protocols.

**Fig. 2.**
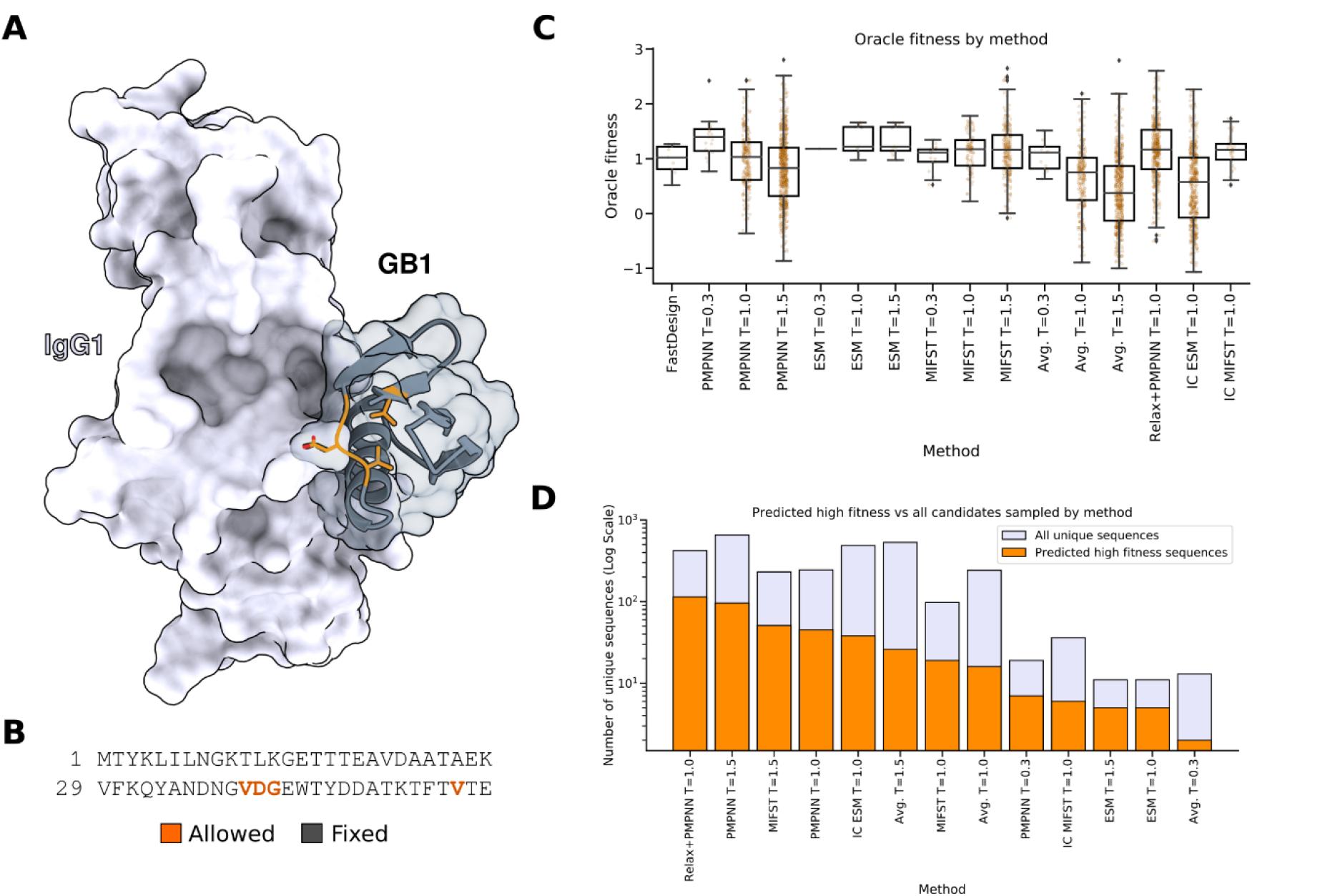
Sampling mutations to improve GB1 fitness. **A)** Structure of GB1 (dark grey) in complex with IgG1 (white) (PDB ID: 1PGA^34^, 1FCC^35^), positions selected for mutation were colored in dark orange. B) Sequence of GB1 with design position in bold and orange. C) Oracle predicted fitness of designed sequences of different methods (wild type equals fitness of one). For each method 1000 trajectories were produced, and all unique sequences were analyzed. D) Bar plot comparing the number of unique sequences sampled by different methods, with orange bars representing predicted high-fitness sequences (greater 1.5) and light purple bars indicating all unique sequences sampled (log scale). PMMPN – ProteinMPNN, ESM – Evolutionary Scale Modeling, MIFST – Masked Inverse Folding with Sequence Transfer, Avg. – Average probability of the three ML models, IC – Iterated Convergence.

As expected, increasing the sampling temperature for amino acid choice led to more diversity (greater number of unique sequences) overall (**Fig. 2 C, D**). For ProteinMPNN and the average of all three models, an increase in temperature led to a decrease in median predicted fitness. In comparison, for both ESM and MIF-ST an increase in temperature did not drastically change the median predicted fitness. While the combination of ProteinMPNN and FastRelax created the most candidates with an oracle predicted fitness greater than 1.5, it also sampled three times as many sequences with a lower predicted fitness (**Fig. 2 D**). In contrast, FastDesign did not create any candidates with a predicted fitness greater than 1.5. Sampling from ESM (T=1.0) resulted only in 11 sequences, however, five of them had a predicted fitness greater than 1.5. Sampling the probabilities from PLMs in an iterative fashion led to different sequence diversities and fitness distributions for both ESM and MIF-ST.

### No method was able to sample a substantial fraction of avGFP variants with improved fluorescence

Another common goal of protein design and engineering is improving or diversifying enzyme activity and selectivity. Therefore, the green fluorescent protein from *Aequorea victoria* (avGFP) was chosen where a dataset compromising 51,715 protein sequences was available^36^. As this dataset covers 81 positions in avGFP (**Fig. 3 A, B**) with an average of 3.7 mutations per sequence, sequences with a maximum of five mutations were generated to ensure reliable oracle prediction, resulting in an average of 3.8 mutations. For this dataset, the fitness was measured fluorescence activity relative to the wild type sequence. Using this dataset, a ridge regression model was trained (spearman correlation of 0.79) and used to predict the fluorescence of generated designs.

**Fig. 3.**
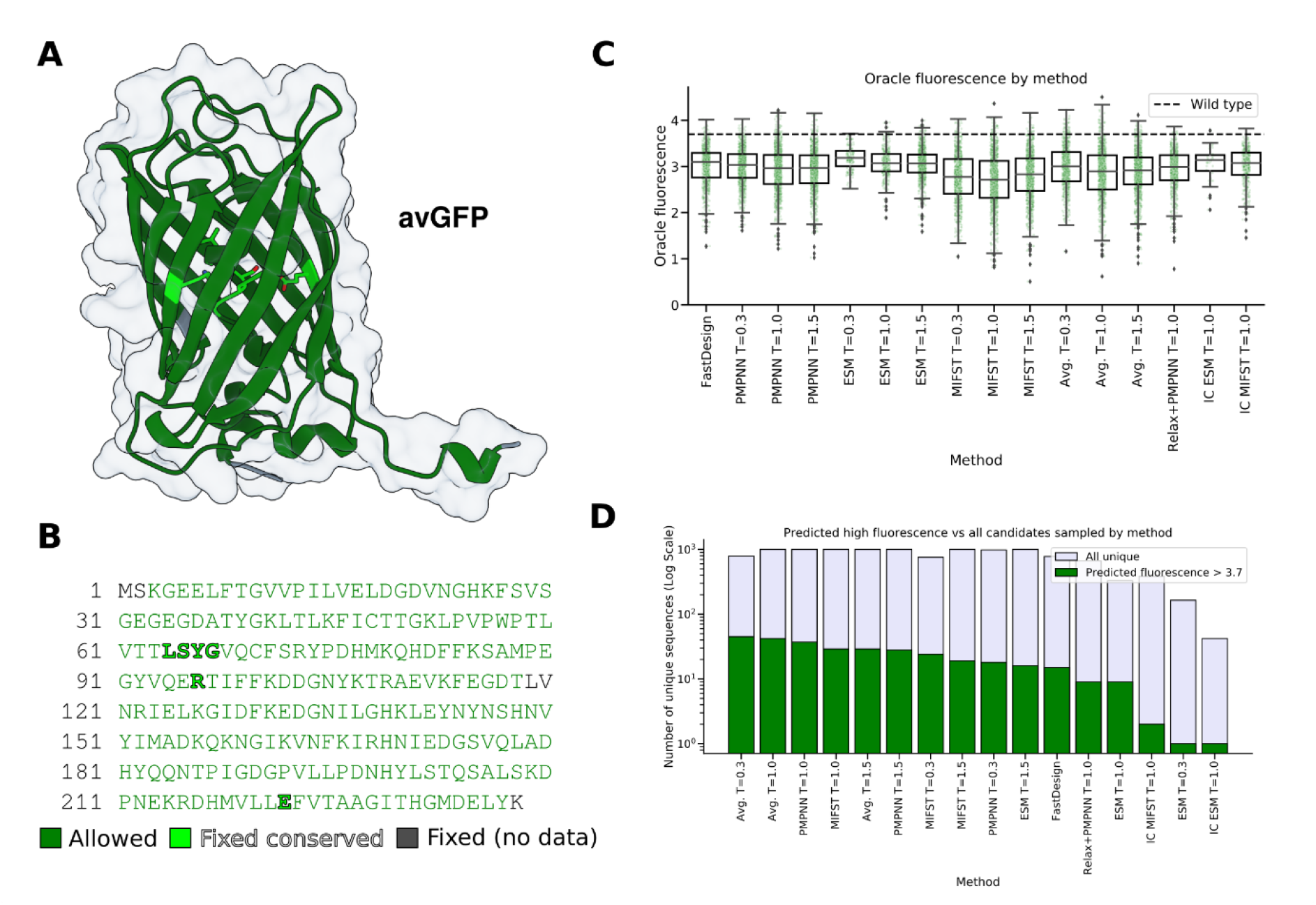
Sampling mutations to improve avGFP fluorescence. **A)** AlphaFold2 predicted structure of avGFP (dark green), positions that were kept fixed in light green (conserved) or grey (no data available). **B)** Sequence of avGFP with design position in dark green. A selected set of highly conserved residues were kept fixed (light green) in addition to positions where no mutagenesis data was available (grey). **C)** Oracle predicted fluorescence of designed sequences of different methods (wild type fluorescence is 3.7). For each method 1000 trajectories were produced, and all unique sequences were analyzed. D) Bar plot comparing the number of unique sequences sampled by different methods, with green bars representing predicted high-fluorescence sequences (greater 3.7) and light purple bars indicating all unique sequences sampled (log scale). PMMPN – ProteinMPNN, ESM – Evolutionary Scale Modeling, MIFST – Masked Inverse Folding with Sequence Transfer, Avg. – Average probability of the three ML models, IC – Iterated Convergence.

Interestingly, none of the design approaches had a median predicted fluorescence greater than the wild type and increased temperatures mainly led to higher diversity of sequences but not to a worse median fitness (**Fig. 3 C**). Sampling from averaged probabilities created the most unique sequences with a fluorescence intensity greater than the wild type, with ProteinMPNN as second best. However, no method had a fraction of these higher fitness candidates larger than 7% (**Fig. 3 D**). Similar to GB1, ESM showed the least number of unique sequences with high fitness compared to other methods. Iteratively sampling from PLM probabilities failed to increase the number of unique sequences with a fluorescence greater than the wild type but increased the diversity of resulting candidates.

### ESM outperforms other methods at diversifying Trastuzumab while maintaining HER2- binding

Next, we selected a case study of generating antibody variants while maintaining binding affinity towards their antigen. We selected a dataset of 38,839 variants of Trastuzumab^37^ (Herceptin), with mutations restricted to ten positions in the variable heavy chain complementarity determining region 3 (CDRH3) (**Fig. 4 A, B**). As for this dataset Trastuzumab variants were classified only into binding and non-binding, we are predicted the binding probability and not the binding affinity, which are not equivalent. To do so, we trained a linear discriminant analysis (LDA) model (0.79 accuracy) to classify binding ability.

**Fig. 4.**
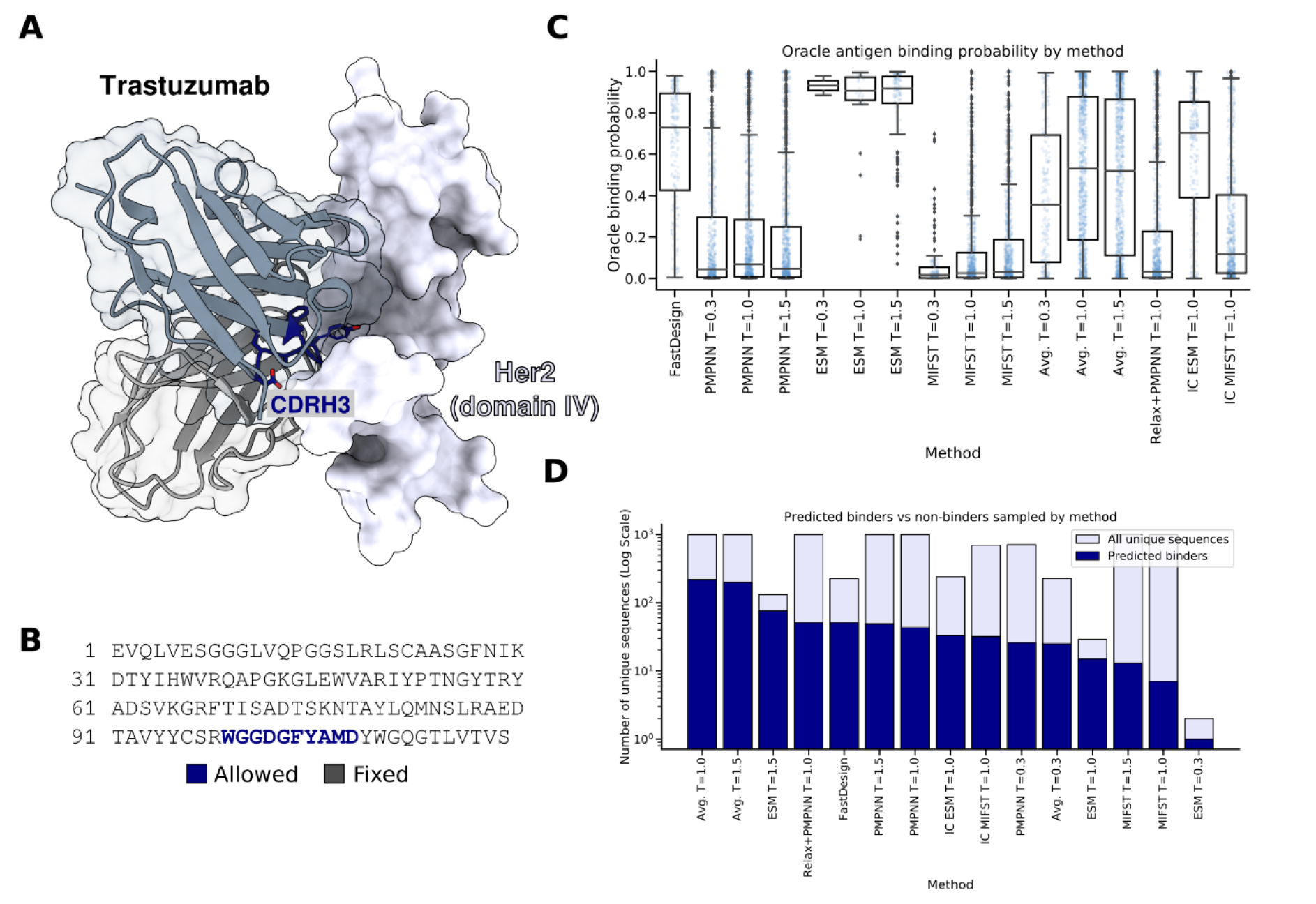
Sampling Trastuzumab variants maintaining HER2-binding. **A)** Structure of trastuzumab (Herceptin) (dark and light grey) in complex with the domain IV of HER2 (white) (PDB ID 1N8Z^38^), positions selected for mutation are colored in dark blue. B) Sequence of the heavy chain of trastuzumab with design position in bold and blue. C) Oracle predicted binding probability of designed sequences of different methods. For each method a thousand trajectories were run, and all unique sequences were analyzed. D) Bar plot comparing the number of unique sequences sampled by different methods, with blue bars representing predicted binders (probability greater 0.9) and light purple bars indicating all unique sequences sampled (log scale). PMMPN – ProteinMPNN, ESM – Evolutionary Scale Modeling, MIFST – Masked Inverse Folding with Sequence Transfer, Avg. – Average probability of the three ML models, IC – Iterated Convergence.

For Trastuzumab, the predicted binding probability of all methods except for (IC-)ESM and FastDesign was below 0.6 (**Fig. 4 C**). Sampling from averaged predictions resulted in the highest number of unique sequences with a predicted binding probability greater than 0.9, but also created five times as many sequences with lower predicted binding probability (**Fig. 4 D**). In contrast, ESM with a sampling temperature of 1.5 resulted in 131 unique sequences of which 76 were predicted to have a binding probability greater than 0.9. Interestingly, while the iterative PLM sampling approach drastically lowered the median predicted binding probability for ESM it improved the results for MIF-ST. Similar to the case of GB1, ESM created a low number of unique sequences of which a greater fraction displayed high fitness values.

### Engineering not only binding affinity but binding specificity remains highly challenging

Lastly, we were interested in increasing the complexity of the design goal, by not only analyzing the binding affinity of generated antibody variants to their antigen, but also whether they display unwanted non-specific (off-target) binding. To do so, we selected a dataset of 10,000 Emibetuzumab variants^39^ which were screened for both antigen and polyspecific reagent binding (**Fig. 5 A, B**). While this dataset is the smallest of the four fitness landscapes used in this study, it offers data on this particularly intriguing dual design objective. Additionally, as no structure of the complex is available, we started from an AlphaFold2^12, 40^ prediction. The resulting structure of the complex is in-line with prior data collected on the epitope^41^. Again, while predicted binding probabilities are not equivalent to binding affinities, they provide insights into the sampling space of different methods used in this study. To classify specificity, we trained two LDA models for predicting either antigen binding (0.87 accuracy) or unspecific binding (0.80 accuracy).

**Fig. 5.**
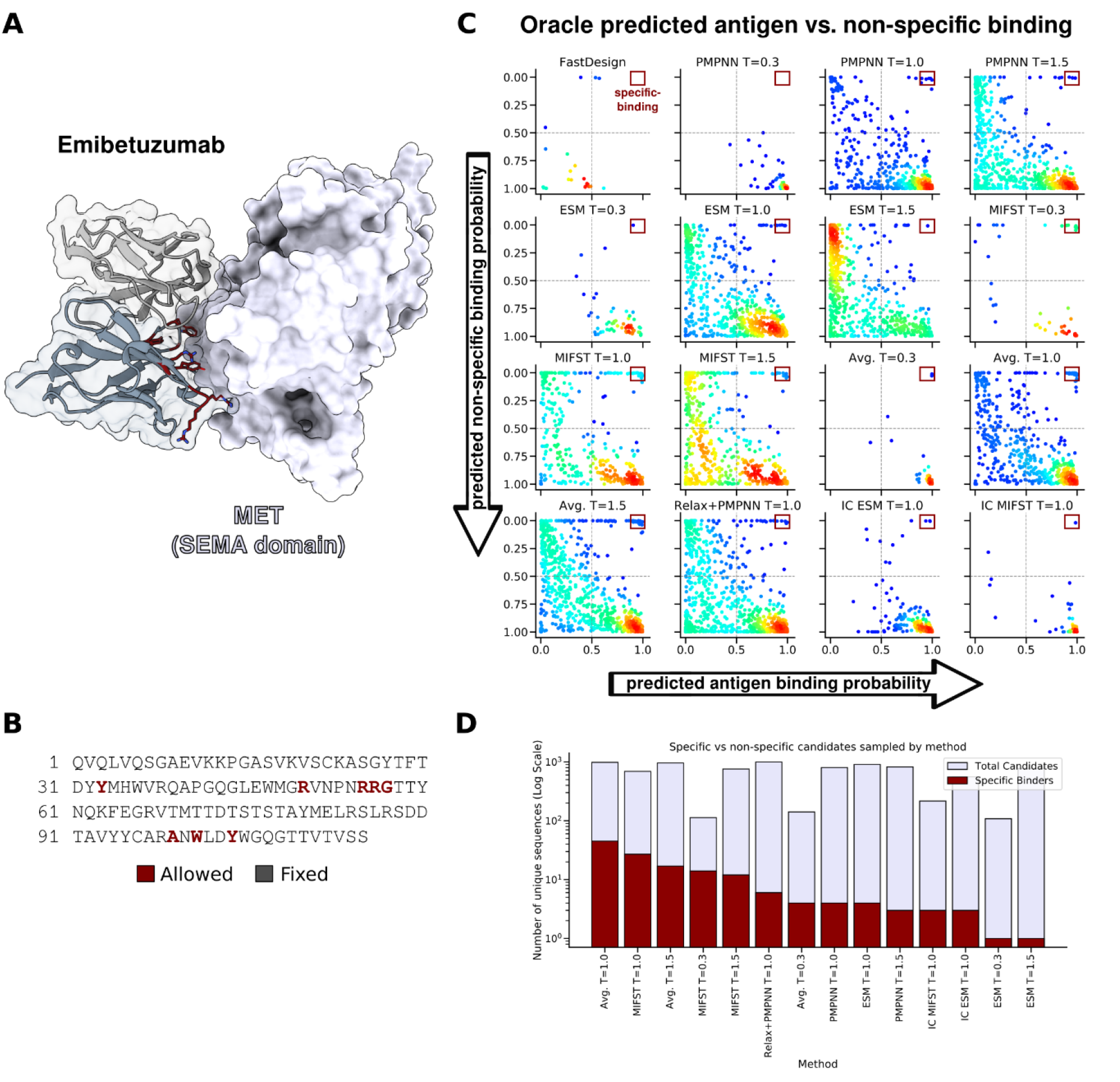
Assessing design approaches on the emibetuzumab dual fitness landscape. **A)** AlphaFold2 predicted structure of emibetuzumab (dark and light grey) in complex with the SEMA domain of MET (white), positions selected for mutation are colored in dark red. B) Sequence of the heavy chain of emibetuzumab with design position in bold and red. C) Oracle predicted antigen binding probability of designed sequences versus oracle predicted non-specific binding probability of different methods. For each method 1000 trajectories were run, and all unique sequences were analyzed. Dots are colored by density ranging from low (blue) to high (red) density. D) Bar plot comparing the number of unique sequences sampled by different methods, with red bars representing predicted specific binders (predicted antigen binding probability greater than 0.9 and predicted non-specific binding probability lower than 0.1) and light purple bars indicating all unique sequences sampled (log scale). PMMPN – ProteinMPNN, ESM – Evolutionary Scale Modeling, MIFST – Masked Inverse Folding with Sequence Transfer, Avg. – Average probability of the three ML models, IC – Iterated Convergence.

For Emibetuzumab, all approaches had a strong tendency to sample sequences that are predicted to bind both the antigen and poly-specific reagents, except FastDesign which only sampled sequences predicted to have low antigen binding probability (**Fig. 5 C**). Sampling from averaged probabilities (T=1.0) created the highest number of predicted specific binders (antigen binding probability greater than 0.9 and non-specific binding probability smaller than 0.1) with 45 unique sequences which were 4.7% of all unique candidates created by this sampling approach (**Fig. 5 D**). Both ESM and ProteinMPNN design resulted in a small (<10) number of predicted specific binders which simultaneously represented a very small (<2%) fraction of sampled sequences. In the case of Emibetuzumab, the iterative approach of sampling from PLMs decreased the overall diversity and in case of MIFST the number of unique candidates with predicted specific binding properties.

### Identifying candidates with highly improved fitness remains challenging

After the successful generation of sequences, ranking and selecting candidates for experimental validation is a crucial step of computational protein design. This ranking is commonly performed by filtering sequences by the score of the model that generated them (e.g. pseudo-perplexity of ML models or total score of Rosetta), as well as biophysical metrics (e.g. shape complementarity, solvent-accessible surface, net charge). Additionally, predicting the structure from the designed sequence and verifying recapitulation of the desired fold can aid in identifying suitable designs, therefore we use AF2 with settings established in prior work^42^. Here, we analyzed the spearman correlation of different scores and metric values from all generated sequences to their oracle predicted fitness scores (**Table 1**). We summarized Emibetuzumab binding properties by calculating the delta for candidates (antigen-specific minus non-specific).

**Table 1.** Spearman correlation between scoring metrics and predicted fitness. The best correlations per protein are marked in bold. NA – Not applicable, AF2 – AlphaFold2, pLDDT- predicted local distance difference test, RMSD – root mean square deviation, pAE – predicted aligned error of the interface. Arrow indicates score direction.

| Spearman p | GB1 fitness | avGFP<br>fluorescence | Trastuzumab<br>binding | Emibetuzumab<br>specific binding |
| --- | --- | --- | --- | --- |
| Total score ↓ | -0.55 | **-0.33** | -0.03 | -0.26 |
| dG separated ↓ | -0.38 | NA | -0.12 | -0.16 |
| Shape<br>complementarity ↑ | -0.02 | NA | -0.09 | 0.11 |
| ESM pseudo-perplexity ↓ | 0.07 | 0.15 | -0.05 | **-0.29** |
| MIF-ST pseudo-perplexity ↓ | **-0.60** | -0.05 | 0.12 | -0.27 |
| PMPNN pseudo-perplexity ↓ | **-0.42** | -0.17 | 0.07 | **-0.29** |
| AF2 pLDDT ↑ | 0.16 | 0.01 | 0.27 | 0.18 |
| AF2 RMSD ↓ | 0.17 | -0.04 | -0.28 | -0.20 |
| AF2 pAE<br>interaction ↓ | 0.04 | NA | **-0.31** | 0.04 |

For the GB1 fitness landscape, MIF-ST pseudo-perplexity showed the strongest correlation (−0.60) to the predicted fitness values, followed by Rosetta total score (−0.55) and ProteinMPNN pseudo-perplexity (−0.42). In the case of Trastuzumab binding probabilities, only very weak correlations were observed except for the AF2 metrics, led by the predicted aligned error (pAE) interaction score. For the predicted GFP fluorescence, only Rosetta total score (−0.33) and ProteinMPNN pseudo-perplexity (−0.17) showed weak correlations. In contrast, all metrics and scores showed weak to moderate correlation for Emibetuzumab specific binding, with ProteinMPNN and ESM leading at −0.29.

Next, we imitated selecting ten candidates for experimental validation by ranking them with the above-described metrics and reported the resulting mean and maximum predicted fitness value of the top ten, as well as the best fitness candidate that could have been found for each fitness landscape in all sampled sequences irrespective of the used method (**Table 2**). As a control, ten candidates were randomly chosen from the sampled pool a thousand times and their predicted fitness values were averaged. Importantly, the random choice was based on the previously sampled candidates and not on random sampling. Emibetuzumab binding properties are again summarized by calculating the delta for candidates (antigen-specific minus non-specific).

**Table 2.** Best design in top ten ranked by different metrics. The best oracle predicted fitness candidates per row are marked in bold and values below random sampling are marked in italic. NA – Not applicable, AF2 – AlphaFold2, pLDDT- predicted local distance difference test, RMSD – root mean square deviation, pAE – predicted aligned error of the interface.

| Mean and best candidate fitness in top ten ranked by | GB1 fitness |  | avGFP fluorescence |  | Trastuzumab binding |  | Emibetuzumab specific binding |  |
| --- | --- | --- | --- | --- | --- | --- | --- | --- |
|  | Mean | Best | Mean | Best | Mean | Best | Mean | Best |
| Total score | 1.22 | *1.58* | 2.80 | 3.88 | **0.70** | 0.92 | *-0.43* | 0.39 |
| dG separated | *0.74* | **2.47** | NA | NA | 0.66 | 0.98 | -0.13 | *0.06* |
| 10% Total score + dG separated | 1.11 | 2.28 | NA | NA | **0.70** | 0.98 | *-0.29* | 0.56 |
| Shape complementarity | *0.68* | *1.73* | NA | NA | *0.23* | 0.90 | *-0.22* | 0.32 |
| ESM pseudo-perplexity | 1.10 | 2.36 | *-4.50* | *-1.22* | 0.35 | 0.97 | -0.02 | 0.06 |
| MIF-ST pseudo-perplexity | 1.20 | *1.56* | *0.20* | 2.28 | 0.29 | *0.46* | **0.13** | **0.97** |
| PMPNN pseudo-perplexity | **1.33** | *1.63* | 2.80 | *3.32* | *0.20* | *0.86* | 0.00 | *0.01* |
| AF2 pLDDT | 1.30 | 2.30 | **2.94** | **3.94** | 0.50 | 0.95 | *-0.04* | *0.15* |
| AF2 RMSD | 1.06 | 1.85 | *2.40* | *3.0* | *0.21* | 0.95 | *-0.06* | *0.15* |
| AF2 pAE interaction | *0.64* | *1.30* | NA | NA | 0.64 | **0.99** | *-0.51* | *-0.07* |
| Random choice | 0.80 | 1.74 | 2.80 | 3.56 | 0.29 | 0.87 | -0.16 | 0.32 |
| Best available | 2.80 |  | 4.79 |  | 1.0 |  | 0.99 |  |

To our surprise, Rosetta total score-based metrics were competitive to ML-based metrics in two out of four cases. Additionally, none of the approaches had the best available predicted fitness candidate in their top ten. For GB1, selecting by each metric would have resulted in at least one candidate with higher fitness, with the dG separated and ESM metrics leading to best selection. Similarly, for Trastuzumab all metrics except for MIF-ST would have selected at least one candidate with high predicted binding probability. In the case of avGFP, only selecting by total score or AF2 pLDDT would have resulted in a candidate with predicted fluorescence greater than the wild type. Most strikingly, all metrics with the exception of MIF-ST perform poorly at ranking Emibetuzumab predicted specific binding.

## Discussion

In this work, we explored the performance of novel self-supervised ML protein engineering methods for sampling and scoring sequences. To assess the newly generated candidates, we trained oracle models on existing large-scale protein fitness datasets to predict the properties of thousands of sequences. We found that data-driven methods excel at limiting the sequence space to non-deleterious mutations compared to “traditional” approaches like Rosetta. When it came to scoring the sampled candidates, however, methods capable of sampling high fitness sequences failed to show a high correlation to predicted fitness values. This dichotomy was especially visible when it came to ranking the sampled candidates, making a selection for experimental validation difficult. Here, models that sampled high fitness candidates and showed a correlation to the predicted fitness values still failed at having these high fitness sequences in their top ten selection. On possible explanation of this observation is the well-known paradigm that’s sampling and scoring are tightly coupled. I.e., if the sampling method is (near) perfect, it outputs results that are indistinguishable by the scoring metric used, in particular since scoring metrics are incomplete, simplified and thus error-affiliated. Scoring the sequences with orthogonal metrics might overcome some of these challenges, in particular if slower but more precise scoring functions can be employed. This orthogonal effect was seen when instead choosing designs based on AF2 metrices for three of our test systems, but not in the more complex design case of Emibetuzumab.

One goal of our study was to find best practices in applying self-supervised ML methods for designing new variants with improved fitness. Both for sampling and scoring, we saw no method constantly outperforming the others. For all four test cases and methods, increasing the diversity through higher temperatures led to broader distribution (more extreme fitness values) while approximately maintaining the median fitness of sampled variants. In our opinion, this gives support to two possible strategies, depending on experimental high-throughput abilities and needed fitness increase. The first strategy is to iteratively sample with very low temperatures and experimentally verify the resulting small number or even single variants, where for each round there is a high likelihood of a reasonable improvement in fitness. The second strategy is to increase the sampling temperature until the resulting candidates match the experimental capabilities and test all of them, which gives the best chances of identifying a variant with greatly improved fitness. In contrast, designing many variants without restraints and then trying to identify the most promising candidates post hoc using *in silico* scores/metrices seems less effective, at least until gaps in scoring functions are filled or supervised ML foundation models become available. The first strategy mentioned closely matches the recent method of creating *de novo* proteins, where many scaffolds are created with e.g. RFdiffusion and then for each only one or two sequences are designed with ProteinMPNN, using a very low sampling temperature^32, 42^. Additionally, the second strategy discussed would be a strong starting point for a ML-guided directed evolution campaign by providing a diverse range of fitness variants as initial dataset. While it is exiting that self-supervised ML models are capable of sampling high fitness sequences, the difficulty in identifying these candidates provides support to these strategies and other approaches which focus on single-point mutations^18, 43^.

Multiple other works have focused on designing proteins with the respective ML models used in this study, for example, to create new antibody variants by sampling single-point mutations from ESM and then combining the most promising in a second round of experiments^18, 43^. Likewise, ProteinMPNN has enabled multiple design and engineering experiments, ranging from hinge-proteins^44^ to solubilizing membrane proteins^45^. While for MIF-ST, to the best of our knowledge, no experimental validation is currently described, the combination of a PLM with structural information is an exciting approach. We initially expected ProteinMPNN to outperform the other methods on all tasks except avGFP fluorescence engineering. We based this on the fact that both ESM and MIF-ST were trained on single chain inputs compared to ProteinMPNN, which could infer with their ability to accurately judge protein-protein interactions, especially in the case of antibodies where less co-evolutionary signal between interaction partners is present. While we only used the to-be engineered protein as input for ESM, we used both interaction partners as input for MIF-ST, inspired by recent work using a similar inverse folding model called ESM-IF by Shanker *et al*.^43^. In contrast to our expectations, both PLMs were able to sample, and to some extent rank, candidates with improved binding properties.

A limitation of our study is the *in silico* approach for validation, and while we informed the oracles with large-scale datasets, the resulting prediction is an approximation. A point to be aware of is the simplicity and additive nature of both the LDA and ridge regression approach used here. Interestingly, previous studies on the Emibetuzumab and Trastuzumab dataset have analyzed both simple and deep learning approaches which are better suited for capturing non-linearity, showing similar performance^37, 39^. Additionally, for GB1 we observed the same results when only taking known sequence-fitness pairs into account or using the oracle model. As previous work has shown that reasoning from single mutations to many is difficult^46^, we restrained the number of mutations for each case to stay close to the average mutations of the corresponding datasets. Nevertheless, the deep self-supervised ML models used for sampling and scoring could have predicted a complex relationship of multiple mutations not picked up by the respective oracle model, resulting in false negatives. However, as the self-supervised ML models were not fine-tuned on the datasets used for oracle training and the number of mutations were limited, a significant amount of these false negatives skewing the overall analysis seems unlikely. Another limitation is the generalization of the case studies to other proteins and systems. In further research the comparison of self-supervised ML methods to case-specific “traditional” protocols could reveal further differences, e.g., evolutionary-informed protocols^47^ for enzyme engineering or antibody tailored methods like RAbD^48^. While we provided a realistic and standardized comparison of self-supervised ML methods, Rosetta also provides various system specific protocols. With the collection of larger and larger datasets, domain specific supervised ML models may be able to directly predict protein fitness.

Taken together, the implemented framework standardizes common tasks involving predicted amino acid probabilities and allows for flexible combinations with existing methods in Rosetta. We showed that while ML models drastically improved sampling of viable sequences, scoring and ranking the created candidates remains challenging. The potential applications of our work include designing proteins ranging from antibodies to enzymes for improved thermostability or functionality and restricting the mutational space towards viable sequences. In conclusion, our work adds novel ways to run, analyze and use predicted amino acid probabilities for protein sequence design and engineering.

## Materials and methods

All methods and protocols are part of this works protocol capture. For further information, please refer to https://github.com/meilerlab/probabilities_design.

### Selection of benchmark datasets

For selecting benchmark case studies, datasets that cover as many variants as possible, while covering different protein attributes were preferred. Therefore, only cases where the measured protein fitness goes beyond “just” protein stability and includes multiple mutations per sequence were chosen. Four different datasets were selected: Protein GB1 compromising ∼150,000 tested mutations of four positions^33, 49^, avGFP with 51,715 test sequences covering 81 residues^36, 49^, a dataset of 38,839 Trastuzumab CDRH3 variants (ten positions)^37^ and 10,000 Emibetuzumab sequences with eight modified positions^39^.

### Oracle model training

For GB1 and avGFP ridge regression was used as oracle model, which has been shown to be a strong baseline, using one-hot encoding and scikit-learn^50^. For Trastuzumab and Emibetuzumab an LDA model was chosen as described by the authors who collected the Emibetuzumab dataset^39^, again using one-hot encoding and scikit-learn^50^. In all cases the full-length sequences were used as input for one-hot encoding.

### Preparation of structure inputs

For GB1, the domain of B1 (PDB ID 1PGA^34^) was aligned with the IgG Fc complex structure (PDB ID 1FCC^35^) and relaxed using the Rosetta relax application^9, 51^, choosing the best scoring candidate from ten trajectories. In the case of avGFP, the structure was predicted using ColabFold^40^ and then relaxed in the same way as for GB1. For Herceptin an available HER2 complex structure (PDB ID 1N8Z^38^) was used and relaxed as before. Lastly, for Emibetuzumab the complex structure with the SEMA domain of MET was predicted using ColabFold^40^ and relaxed as before.

### Sampling protocol details

For the Rosetta baseline, the `FastDesign` mover with the MonomerRelax setting for avGFP and the InterfaceRelax setting for GB1, Trastuzumab and Emibetuzumab was used. For ESM, MIF-ST and ProteinMPNN their respective `PerResidueProbabilitiesMetric` was used to predict and store amino acid probabilities for each protein (called `EsmPerResidueProbabilitiesMetric`, `MIFSTProbabiltiesMetric` and `ProteinMPNNProbabilitiesMetric`). Additionally, the probabilities of all three models were averaged using the `AverageProbabilitiesMetric`. Then the `SampleSequenceFromProbabilities` mover was used to generate sequences from the probabilities and thread them onto the protein structure, using three different sampling temperatures (T=0.3, T=1, T=1.5). For the combination of ProteinMPNN and relax approach, a protocol consisting of three rounds of ProteinMPNN design (using the `ProteinMPNNProbabilitiesMetric` and `SampleSequenceFromProbabilities`) each followed by a relaxation of the structure using the `FastRelax` mover was created. Furthermore, for MIF-ST and ESM the `IteratedConvergenceMover` was executed to sample only one mutation at a time until the pseudo-perplexity score (calculated with the `PseudoPerplexityMetric`) converged. 1000 trajectories were ran for each approach and then all unique sequences were analyzed. Structures were visualized with ChimeraX^52^.

### Scoring and ranking of candidates

All resulting sequences were scored regarding their Rosetta total score (beta update), ML model pseudo-perplexity and, except for avGFP sequences, Rosetta dG separated and shape complementarity. To do so the Rosetta `InterFaceAnalyzerMover,` as well as the PseudoPerplexityMetric which takes as input the respective `PerResidueProbabilitiesMetric` of each model was used. Next, the spearman correlation of each metric with the oracle predicted fitness was calculated using scipy^53^. For the ranking a random baseline was created by choosing ten candidates from the sampled pool a thousand times and averaging the resulting predicted fitness values. Importantly, the random choice and all other metric ranking is based on the previously sampled candidates from all methods and not on random sampling or just the candidates from one specific method. For the AF2 predictions, the structure of designed sequences was predicted using model_4_ptm for avGFP or model_1_ptm for GB1, Emibetuzumab and Trastuzumab. In the case of complex prediction, the structure of all non-mutated chains was given as an initial guess. For all predictions no multiple sequence alignments or additional templates were used.

## Acknowledgment

Computations for this work were done (in part) using resources of the Leipzig University Computing Centre. The authors would like to thank the authors of AF2, ESM, MIF-ST and ProteinMPNN for making their innovative work available.

## Author contributions

Conceptualization: ME CTS. Data Curation: ME. Formal Analysis: ME. Funding Acquisition: CTS JM. Investigation: ME. Methodology: ME RM CTS JM. Project Administration: ME CTS. Resources: CTS JM. Software: ME RM. Supervision: CTS JM. Validation: ME.

Visualization: ME. Writing – Original Draft Preparation: ME CTS JM. Writing – Review & Editing: ME RM CTS JM.

## Data availability

The different benchmark datasets, as well as the sampled sequences and their scores can be found at https://github.com/meilerlab/probabilities_design.

## Code availability

All described features are part of Rosetta which is freely available for academic and non-profit users at http://www.rosettacommons.org/software. A user manual, demos and tutorials can be found at http://www.rosettacommons.org/docs/latest/. The Rosetta source code can be found at https://github.com/RosettaCommons/rosetta/. An overview of features implemented in this paper is given at https://www.rosettacommons.org/docs/latest/scripting_documentation/RosettaScripts/compos <u>ite_protocols/Working-with-PerResidueProbabilitiesMetrics</u> and a specific tutorial can be found at https://meilerlab.org/rosetta-workshop-2023/ (“Tutorial 2: Machine Learning in Rosetta”). The paper specific implemented Rosetta protocols, commands and evaluation code are available at https://github.com/meilerlab/probabilities_design.

