## Supplemental Information for "Self-supervised machine learning methods for protein design improve sampling, but not the identification of high-fitness variants"

### Supplementary Information

### Code availability

All described features are part of Rosetta which is freely available for academic and non-profit users at http://www.rosettacommons.org/software. A user manual, demos and tutorials can be found at http://www.rosettacommons.org/docs/latest/. The Rosetta source code can be found at https://github.com/RosettaCommons/rosetta/. An overview of features implemented in this paper is given at https://www.rosettacommons.org/docs/latest/scripting_documentation/RosettaScripts/composite_protocols/Working-with-PerResidueProbabilitiesMetrics and a specific tutorial can be found at https://meilerlab.org/rosetta-workshop-2023/ ("Tutorial 2: Machine Learning in Rosetta"). The paper specific implemented Rosetta protocols, commands and evaluation code are available at https://github.com/meilerlab/probabilities_design.

### Rosetta Implementation and features

#### Implementation of ML models in Rosetta

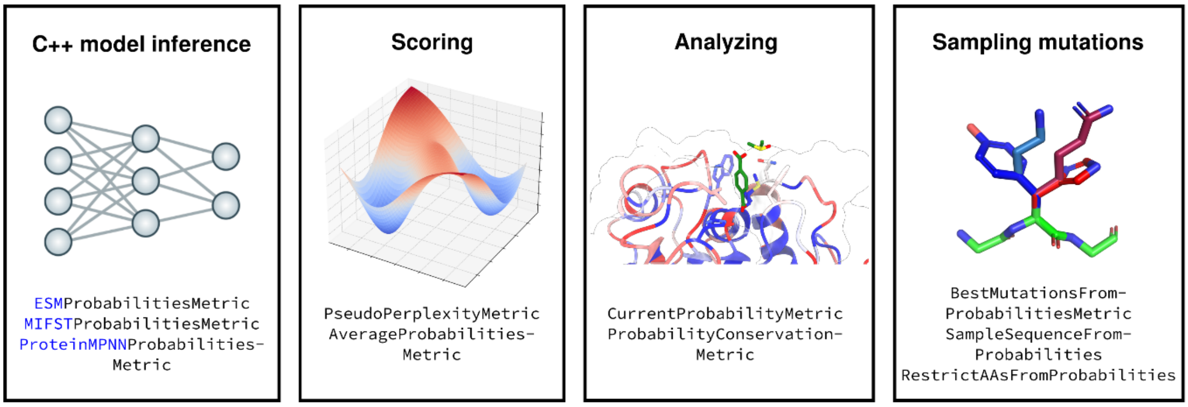


**Fig. S1 ML-support framework in Rosetta.** Implemented RosettaScripts elements include metrics to run different ML models for predicting amino acid probabilities, metrics for analyzing or scoring with the predictions, and movers to directly thread mutations sampled from the predictions on to the structure loaded in Rosetta.

First, we set out to embed current state-of-the-art deep learning models in Rosetta's C++ framework, including feature calculation, inference, and downstream tasks. Recently, new Rosetta builds[1,2] have been created that link against the C/C++ API of TensorFlow[3] and/or LibTorch[4]. This removes the need for Python environment setups, where compatibility between different packages can be non-trivial, and reduces the dependencies to "just" Rosetta and TensorFlow/LibTorch. While Rosetta is a very mature software project that comes with its own complexity and technical debt, the consortium of international academic laboratories known as RosettaCommons have created a rigorous testing server infrastructure[5,6]. This includes testing of various builds for multiple operating systems (including linking against TensorFlow/LibTorch), as well as continuous regression testing of different protocols (including all features described here).

We focus on machine learning approaches that predict amino acid probabilities and are therefore useful for protein design and engineering. To cover a wide set of design objectives, we make use of the protein language model family evolutionary scale modeling (ESM)[7], masked inverse folding with sequence transfer (MIF-ST)[8] and ProteinMPNN[2]. We provide access to running the different models through `RosettaScripts`[9], which is an XML language to define protocols without the need for modifying the underlying C++ code. This allows the user to define different elements and arrange them to a custom protocol without prior scripting or coding experience. The most common objects are either directly changing the provided structure which is defined by the pose object (`Movers`) or evaluating some aspect of it (`SimpleMetrics`). We first created a general `SimpleMetric`[10] class for holding predicted probabilities, called `PerResidueProbabilitiesMetric`. Subsequently, we created a metric for each model that can be used through the `RosettaScripts` framework, where the user can provide a `ResidueSelector` to specify subsets of the protein for prediction. The feature calculation and model inference with specified options stores results in the pose after execution. Through the output standardization, the predictions can be used with the tools described below. While for ProteinMPNN multiple positions can be predicted with one inference call, for ESM and MIF-ST each position is masked and predicted. We provide ways to run the models either on the CPU or GPU.

**Analyzing amino acid probability distributions**

After running a specified model, the predicted amino acid probabilities can be analyzed in various ways. We first provide `RosettaScripts` elements for saving and loading predictions, enabling the split of inference and analysis, as well as loading predictions from an arbitrary source. Furthermore, we established an `AverageProbabilitiesMetric` computing a user defined weighted average of multiple `PerResidueProbabilitiesMetrics`. This averaging is useful to create a prediction ensemble of multiple models or, alternatively, can be used for multi-state design objectives[11] (including negative design). Next, we implemented a `PseudoPerplexityMetric` to assess the likelihood of a sequence given the predicted probabilities, a common score which is defined as the exponentiation of the average negative logarithm of the predicted probabilities. Additionally, we provide ways to distill the predicted probabilities down to either just the probabilities of the amino acids in the current sequence (`CurrentProbabilitiesMetric`) or a score describing whether one particular or many amino acids are seen as likely by the model (`ProbabilityConservationMetric`). These per residue values can subsequently be applied for filtering of design trajectories or for visualization purposes by saving them to the b-factor column of the output file. In some design use cases, e.g., antibody development or enzyme design, the goal is to find the mutations with the most impact on protein stability and/or function without re-designing large parts of the protein.

Inspired by Hie *et al.*[12] we implemented the `BestMutationsFromProbabilitiesMetric`, a metric for finding mutations with the highest delta probability to the current residues. Together with the averaging metric, this allows to quickly ensemble multiple models and generate a list of mutations predicted to have the highest impact. The analysis of predictions can be combined with Rosetta's mutagenesis and design tools.

**Sampling mutations for protein design**

The embedding of ML design models in Rosetta's framework enables the flexible combination with different design protocols. Our first approach is implemented as `SampleSequenceFromProbabilities` mover and focuses on deriving mutations from the predicted probabilities. While sampling mutations for the full sequence of a protein might be useful for e.g., designing a *de novo* protein or the comprehensive re-design of a scaffold, many other common protein engineering tasks seek to maximize impact while minimizing the number of mutations. To cover both use cases, we first rank the to-be-designed positions by the maximum difference of probability to the current sequence, and then rank the amino acids for each position based on their predicted probability. The mover's options allow users to adjust temperature parameters for the selection of both position and amino acid which influence the determinism of their selection, in addition to specifying the desired number of mutations. For example, this interface allows the user to generate full sequences with variability or just identify deterministically combinations of the most impactful mutations.

In addition to sampling directly from predictions, informing Rosetta design with additional information, for example evolutionary signals derived from multiple-sequence alignments[13,14], is a common way of restraining the Rosetta energy function and/or limiting the sequence search space. To achieve the same with predicted probabilities, we implemented a

RestrictAAsFromProbabilities task operation that takes any PerResidueProbabilitiesMetric and allows the user to restrict the designable amino acids to a custom probability cutoff. Furthermore, any PerResidueProbabilitiesMetric can be output as a position-specific-scoring-matrix (PSSM) which can be used with the FavorSequenceProfileMover to restrain the Rosetta energy function with the predicted probabilities during design. For example, we recently showed that the combination of ESM predictions with Rosetta results in more native-like proteins than Rosetta[15].

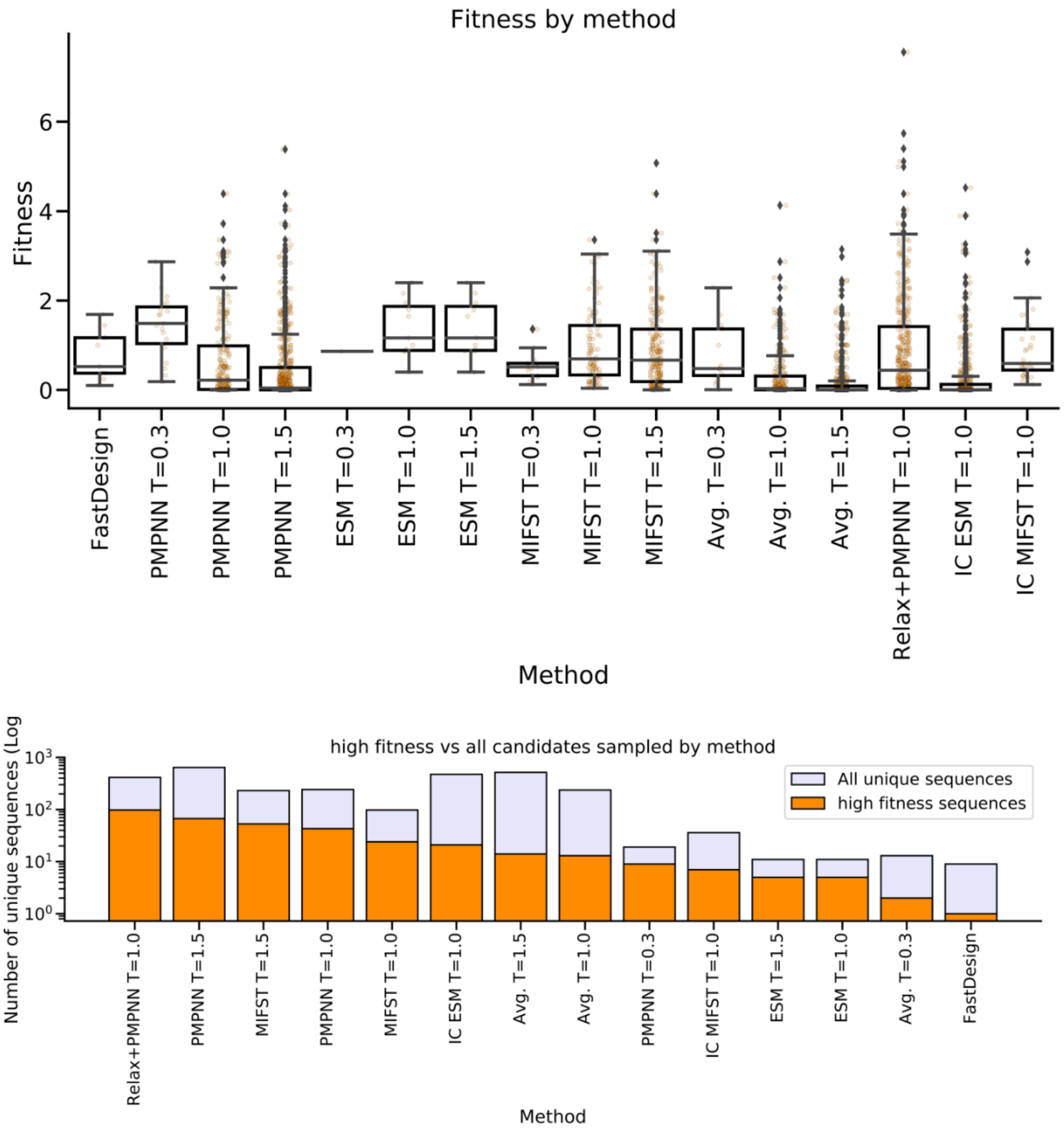


**Supp. Fig. 2 Sampling mutations to improve GB1 fitness.** Top: Oracle predicted fitness of designed sequences of different methods (wild type equals fitness of one). For each method 1000 trajectories were produced, and all unique sequences were analyzed. Bottom: Bar plot comparing the number of unique sequences sampled by different methods, with orange bars representing predicted high-fitness sequences (greater 1.5) and light purple bars indicating all unique sequences sampled (log scale). PMMPN – ProteinMPNN, ESM – Evolutionary Scale Modeling, MIFST – Masked Inverse Folding with Sequence Transfer, Avg. – Average probability of the three ML models, IC – Iterated Convergence.
